# Mitochondrial DNA analysis in ayu (*Plecoglossus altivelis*) raises a urgent conservation need for its southernmost population

**DOI:** 10.1101/2024.01.28.577666

**Authors:** Linh Manh Ha, Hau Duc Tran, Hirohiko Takeshima, Kei’ichiro Iguchi

## Abstract

The central-marginal hypothesis predicts that genetic diversity decreases and genetic differentiation increases toward the species distribution margin. We analyzed the mitochondrial DNA control region of ayu (*Plecoglossus altivelis*) collected from different localities, including central and marginal populations, to test the hypothesis. The results show that the ayu populations at southern marginal habitats (Tien Yen River in Vietnam and Yakugachi River in Amami-oshima Island) displayed low genetic diversity (Haplotype diversity, *H* = 0.52 and 0.67; Nucleotide diversity, *Pi* = 0.01 and 0.006, respectively) and significantly differed from the central populations (*p* < 0.001), consistent with the central-marginal hypothesis. However, ayu populations at the center of their distribution range (Shinano River in Honsyu and Hamoji River in Sado Island) and the northern marginal habitats (Yoichi River in Hokkaido and Aonae River in Okushiri Island) exhibited high genetic diversity (*H* = 1 for all rivers; *Pi* = 0.024–0.029). The genetic differences between the populations at the center and the northern marginal habitats were insignificant (*p* > 0.05). This study showed that in ayu, the central-marginal hypothesis is valid in the southern direction but not in the northern direction. Urgent conservation is needed for the genetically distinct population in the degraded southern habitat with severely low genetic diversity.

## INTRODUCTION

One of the main topics in evolutionary ecology is the study of the characteristics of species populations when they are located at the edges of their geographical range (called marginal populations or peripheral populations, Gaston, 2003). The study of these populations is important for understanding how species respond to climate change, the process of distribution range extension, and speciation (Kirkpatrick and Barton, 1997; Kolzenburg, 2022).

Within a species’s distribution range, environmental conditions typically vary in a gradient of reducing favorableness from central to margins (Mayr 1963; Lawton 1993; Eckert et al. 2008). As a result, populations at the margin of the distribution range often experience less favorable and more stressful environmental conditions compared to their central counterparts (Lawton 1993). This environmental pressure exacerbates genetic drift, resulting in a reduction of genetic variation within these populations (Lesica and Allendorf 1995). Marginal populations are often isolated from the central population and live in patchy habitats (Mayr 1963; Levin 1970; Lesica and Allendorf 1995). As a consequence, marginal populations typically exhibit smaller size, genetic divergence, and lower genetic diversity when compared to central populations (Lesica and Allendorf 1995; Hardie and Hutchings 2010). The widely accepted theory that describes this pattern is the central-marginal hypothesis (CMH) (Mayr 1963; Lawton 1993; Lesica and Allendorf 1995; Eckert et al. 2008).

The ayu, *Plecoglossus altivelis* (Osmeriformes: Plecoglossidae), is an ecologically and commercially important fish. It has a unique life cycle that involves migration between the sea and freshwater during early life stages, and it is classified as an amphidromous fish (McDowall 1992; Iguchi et al. 2005). Adult ayu live and forage in the upper reaches of rivers and streams, primarily feeding on attached algae. During Autumn, sexually mature fish migrate downstream to near the tidal edge, where they spawn and die. Newly hatched larvae drift to coastal areas and spend early life stages in the water near the coast by feeding on zooplankton. In Spring, young fish migrate into the nearby streams or rivers, not necessarily is natal habitat (Kinoshita 1993; Azuma et al. 2003; Iguchi et al. 2006). The species is distributed across Japanese Archipelagoes, the Korean Peninsula, the Chinese coast, and northern Vietnam (Nakabo 2018). Ayu that inhabit in the Ryukyu Islands have been identified as genetically distinct from other populations, leading to their classification as a subspecies known as Ryukyu-ayu (Nishida 1988). In several recent decades, some insular populations in the southern distribution range have gone extinct (in Taiwan; Okinawa Island, Japan) or are in threatened status (in Amami-Oshima Island; Yaku-shima Island) (Nishida et al. 1992; Ministry of the Environment of Japan 2017). This indicates the vulnerability of small populations, particularly those in insular environments and marginal habitats, making them more prone to extinction. Therefore, prioritizing research on the characteristics of populations in marginal habitats is crucial for improved management and conservation strategies.

Based on the predictions of the CMH, we hypothesized that environmental conditions at ayu’s distribution margins would be more severe, hindering dispersal ability and resulting in lower population densities. We further postulated that limited gene flow and reduced population sizes in peripheral populations would lead to (1) lower genetic diversity and (2) stronger localized isolation, causing greater genetic differentiation among marginal populations and those in central areas. By clarifying these issues, we aim to provide crucial information for better management and conservation of the species.

## MATERIALS AND METHODS

### Samples collections

A total of six sample locations were selected across the species distribution range and represented three main distribution portions, including insular and mainland populations at center, northern marginal, and southern marginal populations (Fig. 1). Locality and year of sample collection for each location is as follows: Yoichi River (abbreviation, YOC; Hokkaido, Japan; 140.88 ºE, 43.07 ºN; year of sample collection, 2019); Aonae River (AON; Okushiri Island, Japan; 139.46 ºE, 42.10 ºN; 2005); Hamoji River (HMJ; Sado Island, Japan; 138.34 ºE, 37.98 ºN; 2011); Shinano River (SNN; Niigata Prefecture, Japan; 139.02 ºE, 37.62ºN; 1995); Yakugachi River (YKG; Amami-oshima Island, Japan; 129.40 ºE, 28.25ºN; 2016); Tien Yen River (TIY; Quang Ninh, Vietnam; 107.36ºE, 21.38ºN; 2014, 2015). Fish samples were collected using cast nets, electronic fishing, or gill nets, frozen at -30ºC, or preserved in 99% ethanol before DNA extraction. DNA samples (fin tissue) of fish from Vietnam were brought to Japan for analysis under permission of The Ministry of Natural Resources and Environment, Vietnam (Decision number: 219/QÐ-BTNMT), following the Nagoya Protocol on Access and Benefit-sharing (ABS procedure). The sequences of the same DNA segment from SSN and YKG that were published in previous studies were obtained either by requesting them from the corresponding authors or by retrieving them from DDBJ/International Nucleotide Sequence Database Collection (INSDC)/GenBank. Sources and accession numbers for sequences of each location are as follows: SNN: LC769091–LC769105 (Iguchi et al. 1997); YKG: MZ042268–MZ042311 (Ha et al. 2022).

**Fig. 1.**
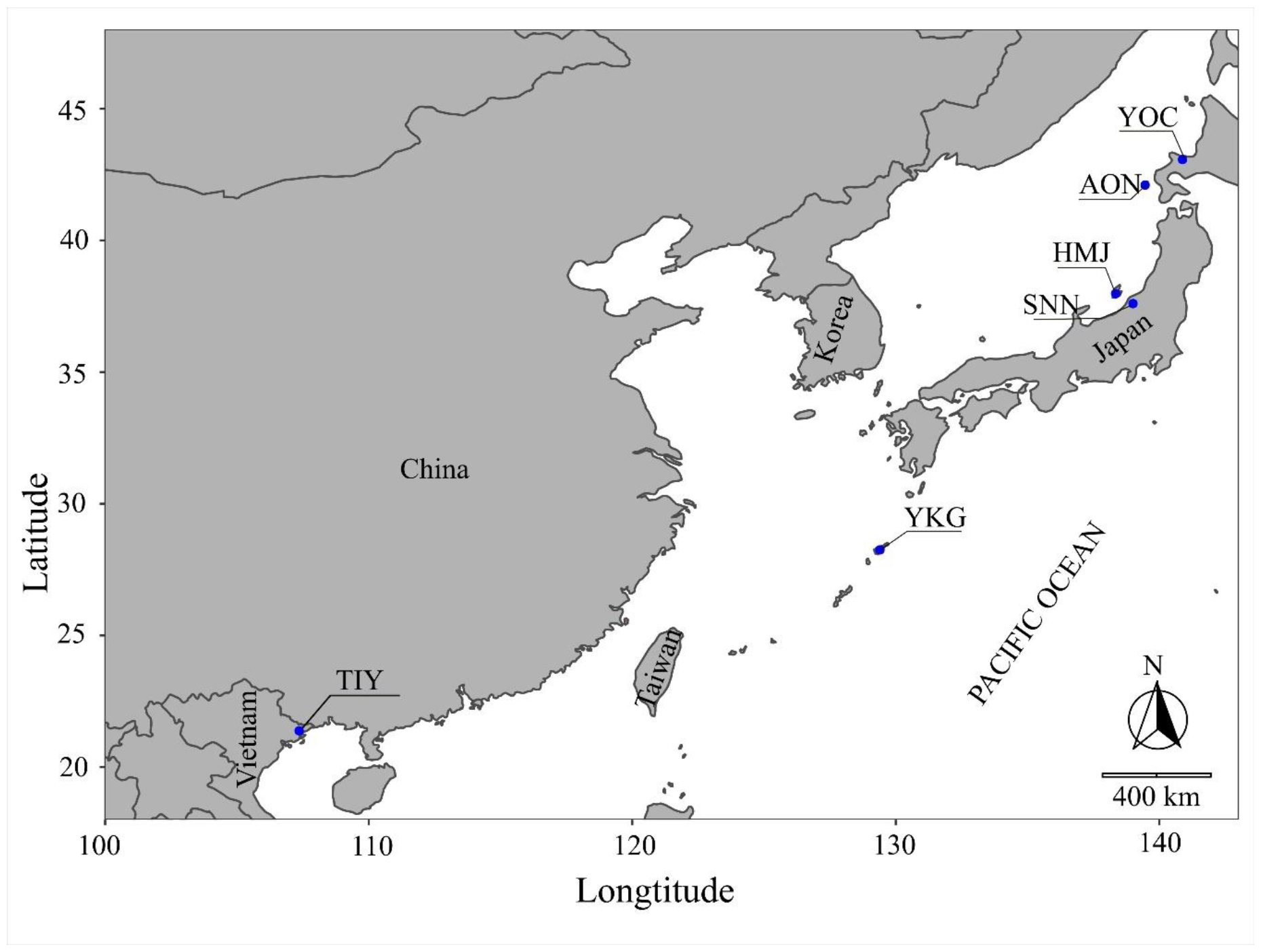
Locations of ayu samples used for mitochondrial DNA analysis; *YOC*, Yoichi River (Hokkaido); *AON*, Aonae River (Okushiri Island); *HMJ*, Hamoji River (Sado Island); *SNN*, Shinano River (Niigata Prefecture); *YKG*, Yakugachi River (Amami-oshima Island); *TIY*, Tien Yen River (Vietnam)

### DNA extraction, amplification, and sequencing

Total DNA was extracted from fin clips or muscle tissues of fish from YOC, HMJ, and TIY using the Gentra Puregene kit following the manufacturer’s protocol (Qiagen, Hilden, Germany). A fragment corresponding to the first half of the mitochondrial DNA control region (D-loop region) was amplified using PCR primers L15923 (5-TTAAAGCATCGGTCTTGTAA-3) and H16498 (5-CCTGAAGTAGGAACCAGATG-3) (Shields and Kocher, 1991), with the standard protocol. PCRs were performed using 7μL reaction volume, containing 3.5 µL GoTaq Green Master Mix 2X, 0.7µL of L15923 primer (5µM), 0.7µL of H16498 primer (5µM), 1µL of DNA template, 1.1µL of Nuclear Free Water. The temperature profile of the PCRs was set at 94ºC for 2 min, followed by 35 cycles of 94ºC for 15s, 50ºC for 15s, and 72ºC for 30s, with a terminal elongation at 72ºC for 7 min. The amplicons were purified using ExoSAP-IT, following the manufacturer’s instructions (Thermo Fisher Scientific Inc., Waltham, Massachusetts, USA). Amplified double-stranded DNA was sequenced using the amplification primer with the BigDye Terminator V3.1 cycle sequencing kit (Thermo Fisher Scientific Inc.) on an automated DNA sequencer ABI PRISM 3100-Avant Genetic Analyzer (Hitachi, Ltd., Tokyo, Japan). All obtained sequences from the specimens of AON, YOC, HMJ, and TIY after trimming to 340bp length are deposited on DNA Data Bank of Japan (DDBJ) under accession numbers LC769020–LC769080.

### Data analysis

Acquired sequences were aligned and trimmed to a common length of 340 bp using MEGA X software (Kumar et al., 2018). Genetic variation in each population was quantified by the number of haplotypes (*H*), haplotype diversity (*h*; Nei 1987), and nucleotide diversity (π; Nei 1987), which were calculated using DnaSP 6.12 (Rozas et al. 2017). Haplotype diversity and nucleotide diversity were compared among populations using the Kruskal-Wallis test, followed by Dunn’s pairwise comparisons in R 4.2.3 (R Core Team 2022). Genetic differentiation among populations (represent by FST values) was revealed by analysis of molecular variance (AMOVA) using ARLEQUIN version 3.5.2 (Excoffier and Lischer 2010).

## RESULTS

A total of 98 sequences of partial fragments of 340-bp from the mitochondrial DNA control region (corresponding to the D-loop region) were acquired from six localities. Analysis of these sequences revealed 73 mutations, with 52 being transitions and 21 transversions. No insertions or deletions were detected. In total, 64 nucleotide positions (18.82%) were found to be polymorphic.

Global analysis of molecular variance (AMOVA) indicated significant genetic differentiation among populations (FST = 0.247, *p* < 0.001). The majority of the variation was found to be among populations (24.70%) and within populations (75.29%). Pairwise FST values and their corresponding *p* values revealed by post-hoc tests among six localities are shown in Table 1. The FST values yielded from comparisons among localities of the center (HMJ and SNN) and those in the northern margin (YOC and AON) were low (ranging from 0 to 0.04), and the differences were not statistically significant after Bonferroni correction. On the other hand, comparisons between fish from localities of the southern distribution range (YKG and TIY) with those of the center and the northern margin all yielded moderate to high and significant FST values (ranging from 0.22 to 0.73; *p* < 0.001).

**Table 1.** Pairwise FST values (above diagonal) and their corresponding *p* values (below diagonal) among populations of ayu (*Plecoglossus altivelis*); *values in italics* indicate statistical significance after Bonferroni correction; *YOC*, Yoichi River (Hokkaido); *AON*, Aonae River (Okushiri Island); *HMJ*, Hamoji River (Sado Island); *SNN*, Shinano River (Nigata Prefecture); *YKG*, Yakugachi River (Amami-oshima Island); *TIY*, Tien Yen River (Vietnam)

| Population | Northern margin |  | Center of Japan |  | Southern margin |  |
| --- | --- | --- | --- | --- | --- | --- |
|  | YOC | AON | HMJ | SNN | YKG | TIY |
| **YOC** | 0 | 0.01358 | 0.00749 | 0.00000 | *0.22843* | *0.41199* |
| **AON** | 0.18018 | 0 | 0.04419 | 0.01952 | *0.31603* | *0.42061* |
| **HMJ** | 0.28829 | 0.03604 | 0 | 0.00000 | *0.26594* | *0.41604* |
| **SNN** | 0.65766 | 0.15315 | 0.42342 | 0 | *0.21511* | *0.39002* |
| **YKG** | *0.00000* | *0.00000* | *0.00000* | *0.00000* | 0 | *0.73242* |
| **TIY** | *0.00000* | *0.00000* | *0.00000* | *0.00000* | *0.00000* | 0 |

Genetic diversity variated among populations (Fig. 2). Haplotype diversity was high in populations of the center (HMJ and SNN; *H* = 15, *h* = 1±0.024) and the northern margin (YOC and AON; *H* = 16, *h* = 1±0.022). Haplotype diversity was significantly lower in populations at the southern margin (*H* = 3, *h* = 0.515±0.081 for YKG in Amami-oshima Island; and *H* = 3, *h* = 0.670±0.082 for TIY in Vietnam; Fig. 2A). Nucleotide diversity showed a similar pattern; it was high for populations in the center (*π* = 0.025±0.002, and *π* = 0.030±0.003 for HMJ and SNN, respectively) and the northern margin (*π* = 0.027±0.002, and *π* = 0.024±0.001 for YOC and AON, respectively). Nucleotide diversity was significantly lower in populations at the southern margin (*π* = 0.010±0.001 for YKG and *π* = 0.00604±0.0006 for TIY; Fig. 2B)

**Fig. 2.**
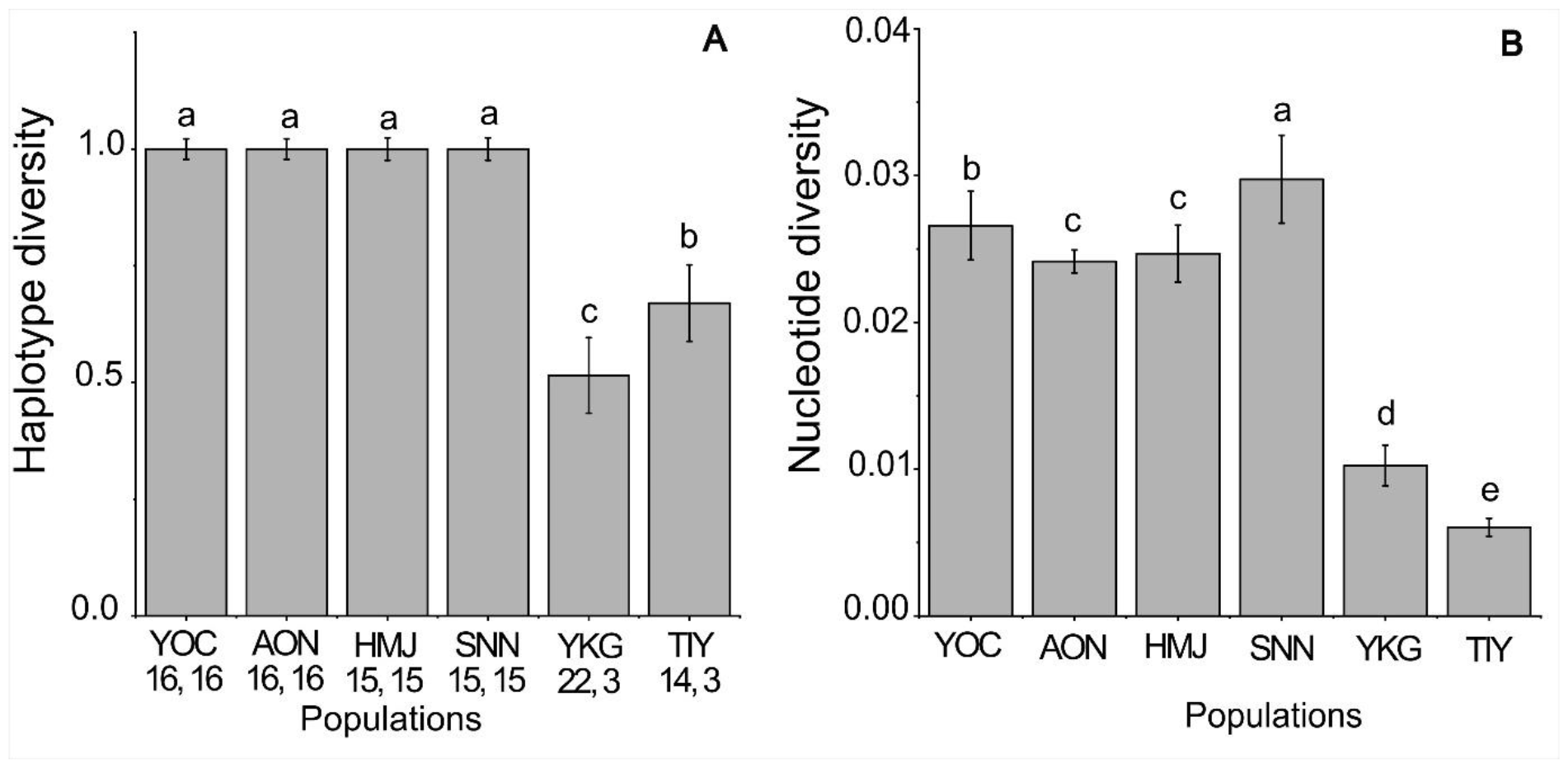
Variation in genetic diversity of ayu populations based on (a) haplotype diversity; (b) nucleotide diversity; *vertical bars* show standard deviation; *differences letters* show statistic differences at *p* < 0.05; *numbers* below the abbreviation of localities indicates sample size and number of haplotypes (*H*), respectively; *YOC*, Yoichi River (Hokkaido); *AON*, Aonae River (Okushiri Island); *HMJ*, Hamoji River (Sado Island); *SNN*, Shinano River (Nigata Prefecture); *YKG*, Yakugachi River (Amami-oshima Island); *TIY*, Tien Yen River (Vietnam)

## DISCUSSIONS

The ayu populations located toward the southern marginal habitats (in Amami-oshima and Vietnam) are less diverse in genetics and more differentiated from those located at the center of the distribution range. The finding for the populations at the southern distribution range is in accordance with the central-marginal hypothesis (Mayr 1963; Eckert et al. 2008; Hardie and Hutchings 2010), which has been confirmed in many other taxa (Lammi et al. 1999; Eckert et al. 2008; Langin et al. 2017; Kitamura et al. 2020; and references herein). The CMH expects that the environmental conditions are less favorable for species at the margin of the distribution range. In Vietnam, the southernmost habitat of ayu, coastal lines are characterized by gradual slopes, expansive river deltas, and extensive mudflats (Eisma 2010). These characteristics make habitats in the southern margin less favorable for ayu, as they typically thrive in small rivers or streams characterized by rapid flows riffles and a substrate composed of cobble and stone. In fact, in Vietnam, there are only two rivers that are located c.a. 60 km apart, where ayu was reported to exist (Tran et al. 2012; Tran et al. 2017; Tran et al. 2018). Such patchy habitats could exacerbate genetic drift and prevent gene flow between populations. As a consequence, ayu populations in the southern distribution area are more differentiated from others in the center, and their genetic diversity is decreased.

In contrast to the southern portion, ayu populations in the northern marginal habitat (Hokkaido and Okushiri Island) are less differentiated, and high genetic diversity is still present. The CMH was not held in the northern portion of the species distribution range. It has been observed in previous studies that the CMH is supported in one direction but rejected in another direction (Böhme et al. 2007; Micheletti and Storfer 2015). In some cases, the center of a species distribution range is skewed to one side, and the CMH is held in the direction to another side (Kitamura et al. 2020). The finding in the present study suggests that even at the margin of the distribution range, environmental conditions in the northernmost habitat are still suitable for ayu to replenish, and gene flow among populations still exists. This is in accordance with previous research suggesting that the ayu populations in the mainland of Japan and adjacent islands constitutes a large population with a metapopulation structure, as indicated by various studies (Iguchi et al. 1999; Iguchi and Nishida 2000; Iguchi and Hirohiko 2006; Kwan et al. 2012). One factor may contribute to suitablility of environemental conditions for ayu is the unique topography of Japan’s coastline. The coastal areas of Japan are rugged and mountainous, with steep cliffs, rocky shorelines, and narrow beaches (Koike 2010). Along the coastal line, there are many short and steep rivers flowing into semi-enclosed bays that provide suitable habitats for ayu that continue from the center to the northern margin (Koike 2010). In other words, ayu populations in the northernmost area are geographically marginal but may not be ecologically marginal.

Furthermore, the high genetic diversity observed in ayu populations residing in the northern marginal habitat can be attributed to the species’ evolutionary origins. Ayu, classified within the Plecoglossidae family as the sole species of the family, is recognized as a relative of Osmeridae in the suborder Salmonoidei, based on shared characteristics (Taniguchi and Ikeda 2009). Osmeridae fishes are primarily found in temperate zones with cooler temperatures. Hence, it is hypothesized that the common ancestor of ayu and Osmeridae fish inhabited temperate habitats. Given ayu’s origin from the temperate zone, colder environments, such as the northernmost habitat in Hokkaido, may still be suitable for their survival.

The applicability of the central-marginal hypothesis in the present study varies between the northern and southern parts of the species distribution, which also could be attributed to the temperature difference of seawater between temperate and subtropical zones. In the temperate zone, ayu spend early life as larval and juvenile at coastal water and can disperse to adjacent rivers before upstream migration (Azuma et al. 2003; Iguchi et al. 2006). This results in intensive gene flow among ayu populations in the main distribution area (Iguchi et al. 2006; Takeshima et al. 2016). In the subtropical zone, high water temperature limits the dispersal ability of fish by lowering the survival rate of fish during marine stages (Takahashi et al. 1999; Kishino and Shinomiya 2005). The period of the marine stage tends to be shorter at a lower latitude (in some cases marine period disappears since fish stay in the brackish water of estuaries) (Kishino and Shinomiya 2005; Murase and Iguchi 2019). Limitation in the dispersal capability of fish results in reduced population size and gene flow among adjacent rivers at low latitude areas and, in turn, may increase differentiation among populations. Therefore, it is explicable when ayu populations in the southern marginal habitats showed higher differentiation and lower genetic diversity compared to those at the center.

The results of the present study also raise concerns about the ayu population in Vietnam, which is genetically different from other populations and has critically low levels of genetic diversity (equal to the level of endangered subspecies in Amami-oshima Island). The habitat for ayu in this southernmost area is limited, patchily, and isolated as is on an island. Environments continually change, and global warming adds stress to this marginal population, but low genetic variation may not allow the population to evolve (Frankel and Soulé 1981), making it more prone to extinction. This suggests an urgent need for conservation measures to protect this ecologically and commercially important species, particularly in the southernmost habitat, which is important for the evolutionary process (Lesica and Allendorf 1995).

## Author contribution statement

LM Ha: conceptualization; writing – original draft, methodology, data curation, data analysis. HD Tran: writing – review & editing, sample collection. H Takeshima: methodology, writing– review & editing, data curation. K Iguchi: resources, methodology, writing – review & editing, conceptualization, supervision.

## DECLARATION

We (the authors) hereby declare that we have no conflicts of interest that could influence the objectivity, integrity, or impartiality of the research presented in this article.

During the preparation of this work the author(s) used Chat-GPT in order to improve readability of the text. After using this tool/service, the author(s) reviewed and edited the content as needed and take(s) full responsibility for the content of the publication.

## ACKNOWLEDGEMENT

We would like to convey our heartfelt gratitude to Prof. Dr. Masaki Nagae of Nagasaki University for generously providing access to his laboratory facilities for DNA analysis. We also extend our sincere appreciation to Prof. Dr. Yoshitaka Sakakura, also of Nagasaki University, for his invaluable comments on the manuscript.

